# Locus-coeruleus norepinephrine activity gates sensory-evoked awakenings from sleep

**DOI:** 10.1101/539502

**Authors:** Hanna Hayat, Noa Regev, Noa Matosevich, Anna Sales, Elena Paredes-Rodriguez, Aaron J Krom, Lottem Bergman, Yong Li, Marina Lavigne, Eric J. Kremer, Ofer Yizhar, Anthony E Pickering, Yuval Nir

**Affiliations:** Department Physiology and Pharmacology, Sackler School of Medicine, Tel Aviv University, Israel; Sagol School of Neuroscience, Tel Aviv University, Israel; School of Physiology, Pharmacology and Neuroscience, University of Bristol, Bristol, BS8 1TD, UK; Department of Pharmacology, Faculty of Medicine and Nursing, University of the Basque Country (UPV/EHU), 48940, Leioa, Spain; Neurodegenerative Diseases Group, Biocruces-Bizkaia Health Research Institute, 48903, Barakaldo, Spain; Department of Anesthesiology and Critical Care Medicine, Hadassah-Hebrew University Medical Center, Hebrew University-Hadassah School of Medicine, Jerusalem, Israel; Institute de Génétique Moléculaire de Montpellier, University of Montpellier, CNRS, Montpellier, France; Department of Neurobiology, Weizmann Institute of Science, Israel; Department of Anaesthesia, University Hospitals Bristol, Bristol, BS2 8HW, United Kingdom; Functional Neurophysiology and Sleep Research Lab, Tel-Aviv Sourasky Medical Center, Tel Aviv 64239, Israel

**Keywords:** Auditory, LC, noradrenaline, arousal threshold, NREM, REM, optogenetics

## Abstract

A defining feature of sleep is reduced responsiveness to external stimuli, but the mechanisms gating sensory-evoked arousal remain unclear. We hypothesized that reduced locus-coeruleus norepinephrine (LC-NE) activity during sleep mediates unresponsiveness, and its action promotes sensory-evoked awakenings. We tested this using electrophysiological, behavioral, pharmacological, and optogenetic techniques alongside auditory stimulation in freely behaving rats. We found that systemic reduction of NE signaling lowered probability of sound-evoked awakenings (SEAs). The level of tonic LC activity during sleep anticipated SEAs. Optogenetic LC activation promoted arousal as evident in sleep-wake transitions, EEG desynchronization, and pupil dilation. Importantly, liminal LC excitation before sound presentation increased SEA probability. Optogenetic LC silencing using a soma-targeted anion-conducting channelrhodopsin (stGtACR2) suppressed LC spiking and constricted pupils. Brief periods of LC opto-silencing reduced the probability of SEAs. Thus, LC-NE activity determines the likelihood of sensory-evoked awakenings and its reduction during sleep constitutes a key factor mediating behavioral unresponsiveness.

## Introduction

Sleep is characterized by a reversible disconnection from the environment that entails reduced responsiveness to external stimuli. An elevated arousal threshold is a central feature of sleep that is present in all mammals and constitutes the main criterion by which sleep is defined in invertebrates lacking cortical EEG such as reptiles, nematodes, flies and fish, and even in jellyfish without a central nervous system ^1–4^. Thus, understanding how sleep is maintained in the face of external sensory events, and what determines sensory-evoked awakening is an open question. Naturally, the intensity of sensory stimuli affects awakening probability ^5^, but other factors such as behavioral relevance also play a role. For example, vocalizing a person’s name is more likely to produce awakening compared to a meaningless stimulus of equal intensity ^6, 7^. Apart from the properties of sensory stimuli, the internal state also plays a role. For example, arousal thresholds depend on sleep stage (NREM sleep vs. REM sleep), sleep duration, and slow wave activity ^5^. In addition, arousal thresholds vary substantially between individuals and with age ^8, 9^: while some individuals experience hyper-arousal (e.g. chronically, or during transient anxiety) and wake up frequently from weak stimuli, others sleep deeply and only high-intensity stimuli lead to awakenings. What determines sensory-evoked awakening and the factors underlying variability across individuals, age, and throughout sleep remain largely unknown.

Wakefulness is supported by a number of subcortical wake-promoting neuromodulatory systems ^10^. Following the identification of the ascending reticular activating system ^11^, a more detailed parcellation of wake-promoting systems followed. Multiple studies established that levels of norepinephrine (NE), serotonin, histamine and hypocretin (orexin) are high during wakefulness and low during both NREM and REM sleep ^12–16^ whereas acetylcholine levels and dopaminergic levels (in the ventral tegmental area and in the ventral periaqueductal gray matter) are high during both wakefulness and REM sleep and decrease during NREM sleep ^17–19^. Recent studies established a causal relationship between neuromodulatory activity and awakenings. For example, optogenetic activation of hypocretin neurons in the posterior hypothalamus ^20^, noradrenergic neurons in the locus-coeruleus ^21, 22^ and dopaminergic neurons in the dorsal raphe nuclei ^23^ increases the probability of sleep-to-wake transitions. Activating GABAergic neurons from the lateral hypothalamus, dopaminergic neurons (in the VTA and ventromedial thalamic nucleus) induces rapid awakening from NREM, but not REM sleep ^18,24,25^. However, these experiments focused on internally triggered sleep-to-wake transitions and it remains unclear whether the same systems also mediate sensory-evoked awakenings.

We hypothesized that the LC-NE system plays a key role in mediating sensory-evoked arousals, and that its reduced activity throughout sleep contributes to unresponsiveness. Several lines of evidence support this proposition: first, LC activity was found to be high during wakefulness, low during NREM sleep, silent during REM sleep and to increase prior to sleep-to-wake transitions ^12^. Second, optogenetic LC activation leads to immediate awakening within few seconds in mice ^21^ and rats ^22^, whereas activation of hypocretin neurons leads to an awakening with latencies around 25s (Adamantidis et al. 2007), and those are likely mediated via LC projections ^26^. Third, LC-NE activity is implicated in orienting responses ^27^, in responding to salient behaviorally-relevant stimuli ^28, 29^, and is involved in stress and pain processing that affects arousals and awakenings ^30, 31^. Fourth, LC-NE modulates the signal-to-noise ratio (SNR) in sensory systems ^32^. Fifth, a higher arousal threshold is found in NE-deficient mice after sleep deprivation ^33^, and low NE levels during wakefulness are associated with electrophysiological markers of disengagement from the environment such as high-voltage spindles ^34, 35^. Given these considerations, we set out to test the causal role of LC-NE activity in mediating sensory disconnection during sleep.

We performed a series of experiments in rats, focusing on auditory stimulation during natural sleep where we controlled sensory stimulation with high temporal resolution. After establishing an experimental system to quantify auditory arousal thresholds from natural sleep, we pharmacologically manipulated NE signaling, finding that lower NE signaling reduced awakening probability. Next, we recorded LC spiking activity during natural sleep and found that baseline tonic LC activity was higher before sound that produced awakenings. To test the causal role of LC-NE activity in more detail, we established selective and efficient optogenetic control over LC activity, including the use of a new vector to silence the LC. Both optogenetic excitation and silencing reliably controlled LC activity and bi-directionally modulated pupil size. We then delivered sounds during natural sleep in conjunction with either liminal LC excitation or with LC silencing, finding that LC-NE activity is sufficient (and necessary) to modulates the probability of awakening in response to sound.

## Results

### Attenuation of NE signaling decreases the probability of sound-evoked awakening (SEA)

We first established and validated an experimental protocol to reliably quantify auditory arousal threshold in freely sleeping rats. We continuously monitored the vigilance state via EEG, EMG, and video as animals spontaneously switched between wakefulness and sleep (Fig. 1a). Tone pips were intermittingly presented with long inter-stimulus intervals, and for each trial we determined if the stimulus led to an awakening or whether sleep was maintained (Fig. 1b, Supp. Fig. 3 & Methods). The probability of SEA monotonically increased with sound intensity levels (Fig. 1c). For example, awakening probability significantly increased by 27 ± 4% (NREM sleep) and 49 ± 10% (REM sleep) as sound intensity increased from 60 dB to 90 dB SPL (n=4 rats), and significantly increased further by 61 ± 1% (NREM sleep) and 59 ± 9% (REM sleep) as sound intensity increased from 80 dB to 90 dB (n=2 rats). The increase in awakening probability as a function of sound intensity established that arousal threshold can be quantified in a sensitive and reliable manner using this approach. In addition, we find higher EEG slow wave activity (SWA, 0.5-4 Hz) in pre-stimulus baseline (2s before sound onset) in trials not followed by behavioral awakenings (Supp. Fig. 3d), as previously shown ^5^.

**Figure 1.**
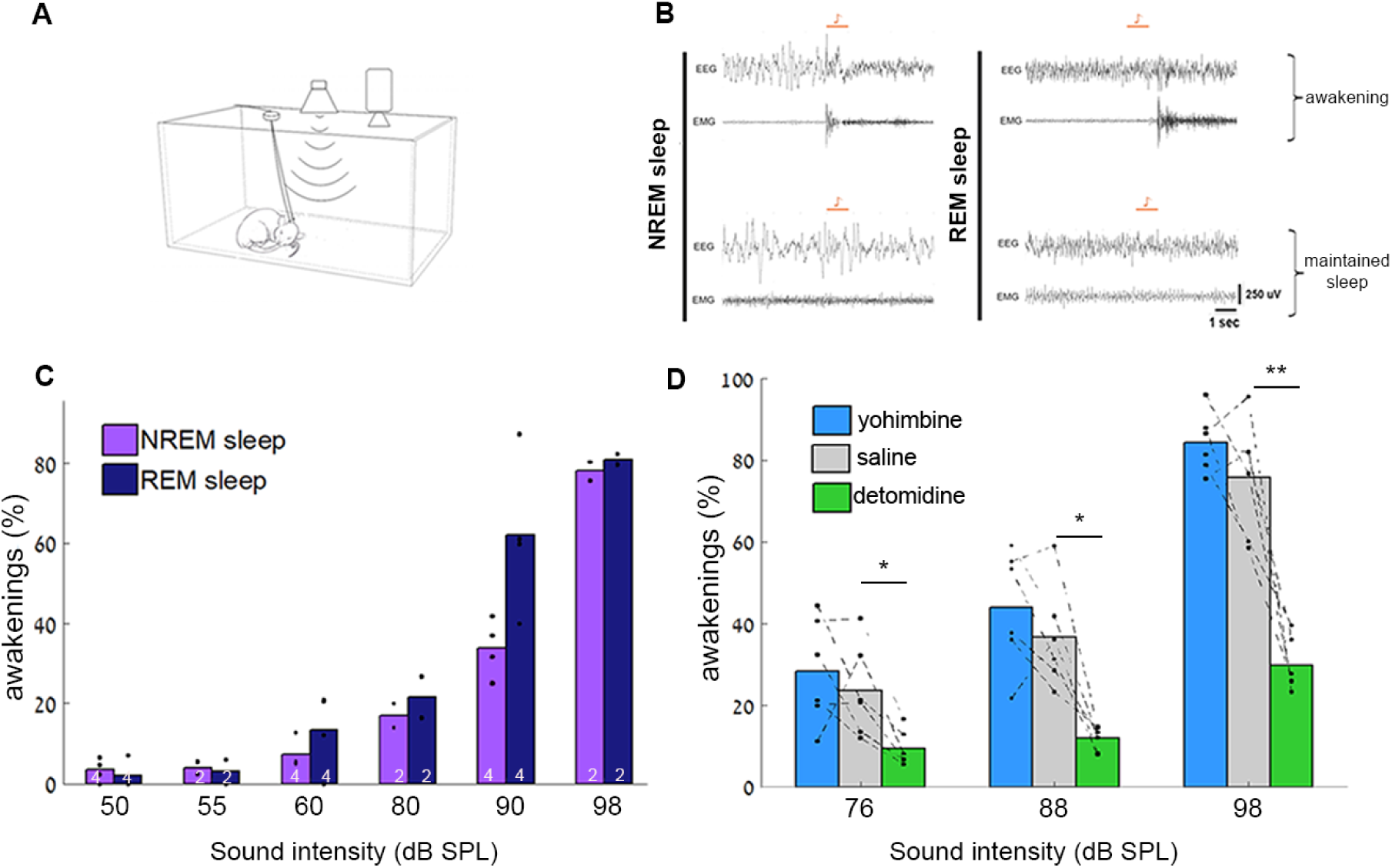
Lower NE signaling decreases the probability of sound-evoked awakenings from NREM sleep. (A) Schematic of experimental setup for rat arousal threshold experiments. Sounds were delivered intermittently from speaker on top while animals continuously monitored with EEG, EMG, and video. (B) Representative EEG and EMG traces showing immediate awakenings (top) vs. maintained sleep (bottom) after auditory stimulation (4kHz pure tone, 1s duration, orange bars) in NREM sleep (left) and REM sleep (right). (C) Probability of awakenings (%) as a function of sound intensity in NREM sleep (purple) and REM sleep (dark blue). White numbers on bars mark number of animals contributing to each result. Average of n=2.75 (4.5) experiments per rat for n=4 (2) rats. Two-way repeated measure (RM) ANOVA reveals significant interaction between sleep stage and sound intensity (F=5.07, p=0.018, n=4 rats), and main effects of sound intensity (p=4.15×10^−8^ for n=4 rats, p=6.18×10^−7^ for n=2 rats) and sleep stage (p=0.01, n=4 rats) (D) Probability of awakenings (%) as a function of sound intensity in NREM sleep following administration of detomidine (α2-agonist, lower NE, green), yohimbine (α2-antagonist, higher NE, blue), or saline (gray). Note that lower NE decreases awakening probability. Two-way RM ANOVA followed by post-hoc t-tests corrected with False Discovery Rate (FDR) *p<0.05, **p<0.01 in n=6 rats.

We then evaluated SEA probability after NE drug interventions. We injected either detomidine (α2 agonist, to decrease NE signaling, 1 mg/kg), yohimbine (α2 antagonist, to increase NE signaling, 1 mg/kg) or saline (control) intraperitoneally (i.p.). We first evaluated the effects of modulating NE signaling on sleep architecture and locomotor activity (Supp. Fig. 1). Detomidine significantly increased the time spent in NREM sleep at the expense of wakefulness and REM sleep (20.2 ± 1.1% increase in NREM sleep, n=6 rats, Supp. Fig. 1b). Yohimbine did not significantly affect sleep architecture but did increase locomotor activity during wakefulness (Supp. Fig. 1d). Next, we evaluated SEAs after NE drug injections. Detomidine significantly decreased the probability of SEA during NREM sleep (14.1 ± 3.4%, 24.8 ± 5.0% and 46.1 ± 7.3% decrease for 76, 88 and 98 dB SPL sound levels, respectively; n = 6 rats Fig. 1d and Supp. Fig. 1a). Yohimbine did not affect the probability of SEA. Although this systemic pharmacological manipulation has some caveats given the widespread actions of α2-adrenoceptor agonists, nonetheless this supported the suggestion that attenuating NE signaling decreases the probability of SEA.

### Baseline tonic LC activity reflects probability of SEA from NREM sleep

To determine if ongoing changes in LC activity predict SEAs during sleep, we recorded the spiking of LC neurons (n = 9 units (8 multi-units/1 single-unit), from one rat) using chronically implanted, drivable silicon probes in freely moving animals. LC neurons were identified by their typical action potential waveforms and their “bi-phasic” responses to toe pinch ^30, 36^ (Fig. 2a). We found that mean spike discharge rates were highest during active wakefulness, lower during quiet wakefulness, and lowest in sleep (Fig. 2d), as previously reported ^12^. The lowest firing rates were observed during NREM-to-REM transitions, when sleep spindle activity is maximal ^37–39^. LC neurons reliably responded to auditory stimulation by firing brief bursts shortly after sound onset (Fig. 2b,e). Dividing auditory trials during NREM sleep to those that ultimately led to an awakening or not (Fig. 2c) we found that LC spiking activity before sound onset was significantly higher before trials that led to awakening (1.99 ± 0.49 Hz) vs. maintained sleep (1.41 ± 0.39 Hz) (Fig. 2c). The phasic response (0-100ms after sound onset) was not linked with SEA probability during NREM sleep (p = 0.959, paired t-test, n = 9) and showed an opposite trend relative to tonic baseline firing (phasic response: active wake<quiet wake< NREM sleep, n = 9) (Fig 2g). We also noted that the higher the spontaneous activity before the sound, the lower was the phasic response to the sound (n = 9, Fig. 2h).

**Figure 2.**
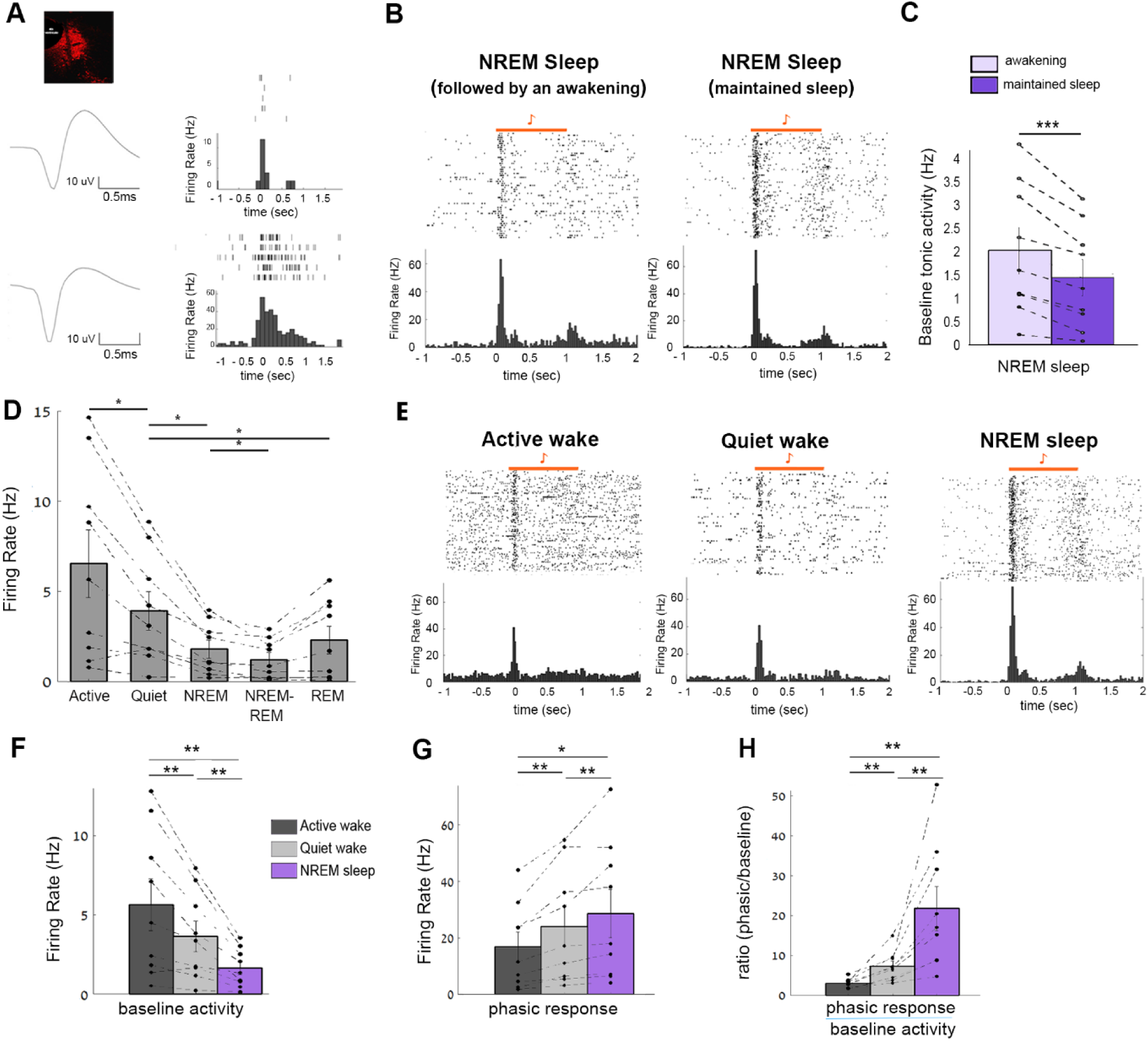
Baseline tonic LC activity is higher prior to sound-evoked awakenings from NREM sleep. (A) Top, image of electrode track targeting the LC superimposed with TH labeling (red). Bottom, two representative LC units recorded during the experiment. Left, action potential waveforms. Right, corresponding raster plot and PSTH in response to toe pinch under light anesthesia. (B) Representative auditory-evoked multi-unit (MU) firing (raster and PSTH) during NREM sleep. Left, trials followed by awakening; Right, trials followed by maintained sleep. Horizontal orange bar shows 1s tone stimulus (C) Quantification of baseline LC activity during NREM sleep in trials preceding awakening (light purple) vs. maintained sleep (dark purple). Data represent mean ± SEM (***p<0.001, paired t-test, n=9 units). Note significantly higher baseline activity prior to awakenings (41% increase compared to maintained sleep). (D) Average firing rates of LC units (8 MUs/1 SU) across different sleep/wake states, one-way RM ANOVA followed by post-hoc FDR-corrected t-test (*p<0.05). (E) Representative auditory-evoked LC MU firing (raster and PSTH) during active wake, quiet wake, and NREM. (F) LC baseline activity 2s before sound onset as a function of vigilance state (G) Phasic LC response to sound (0-100ms after sound onset) as a function of vigilance state and (H) Ratio between phasic response and baseline activity as a function of vigilance state (F-H) Mean ± SEM across n = 9 units, one-way RM ANOVA followed by post-hoc FDR-corrected t-test, *p < 0.05, **p < 0.01.

### Optogenetic LC excitation elicits behavioral, electrophysiological, and pupillary signs of arousal

To achieve cell-type specific optogenetic control over LC activity, we unilaterally transduced LC neurons by injecting CAV2-PRS-ChR2-mCherry (Fig. 3a). Transgene expression was specific and effective as 83.1% of the TH^+^ neurons expressed mCherry (Fig. 3b). Next, we verified optical activation of LC neurons in vivo under light anesthesia. We implanted a 16-channel optrode and recorded LC firing activity in neurons in the dorsal pons, which exhibited characteristic wide action potentials and responded to both toe pinch (Fig. 3c) and laser stimulation (Fig. 3d,e,f,g). Upon laser illumination, we found reliable activation of LC neurons that showed a clear relationship between firing rate and illumination parameters including light intensity, pulse duration, and stimulation frequency. Although a subset of LC neurons may not respond to toe pinch ^30,40,41^, in our hands neurons that responded to laser stimulation showed toe pinch-evoked firing which had a typical biphasic response to contralateral and ipsilateral pinch ^42, 43^(Fig. 3c).

**Figure 3.**
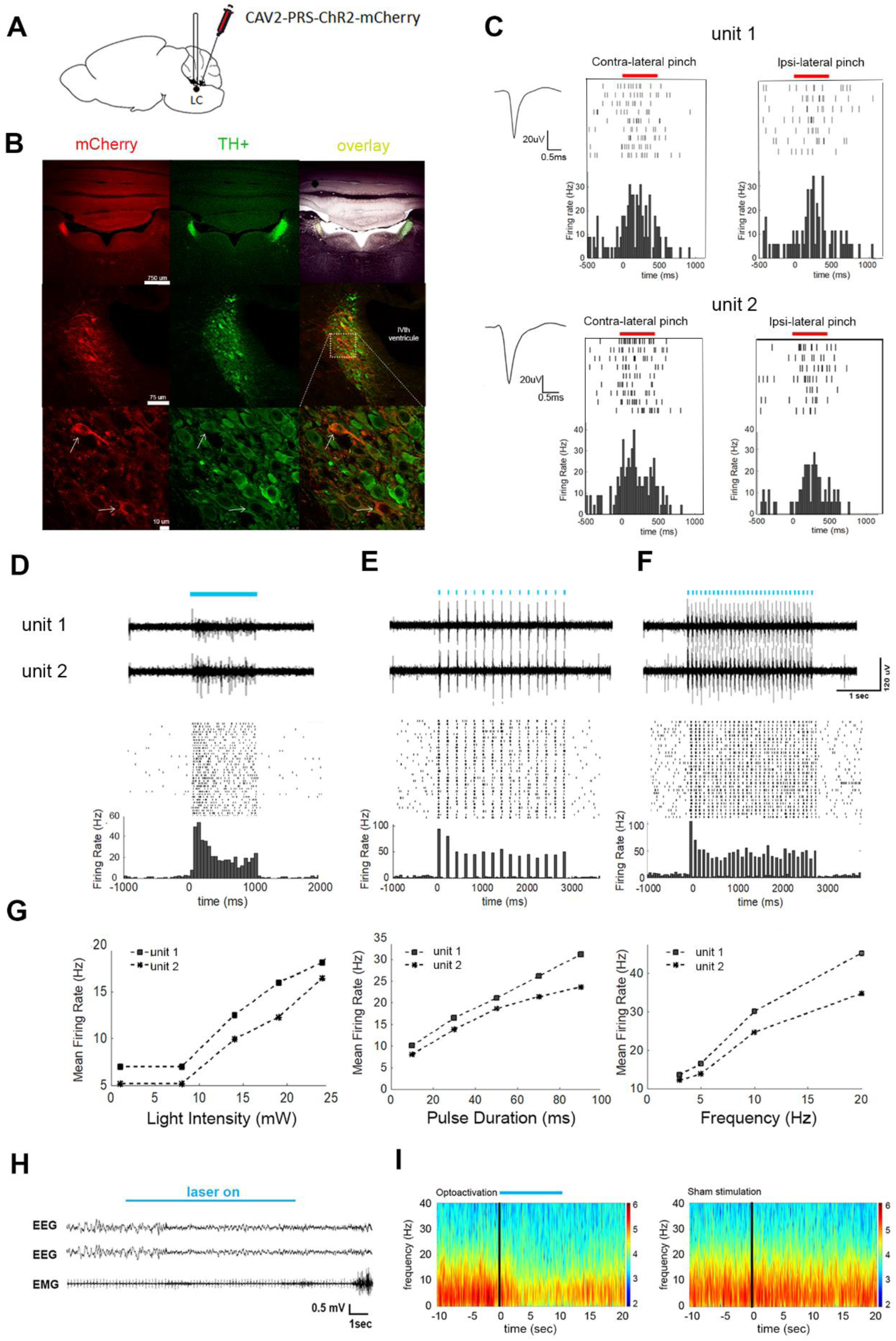
Specific and effective optogenetic excitation of LC neurons. (A) Schematic of unilateral LC injection and optic fiber implantation. (B) Specific expression of ChR2-mCherry in LC neurons: representative coronal images showing CAV2-PRS-ChR2-mCherry expression (left column, red), labeling of TH^+^ neurons (middle column, green), and their overlay (right column, yellow). Top images, global expression in coronal sections; middle row, expression around the LC; bottom row, magnified images of cells in the boxed area marked in middle row. Arrows mark neurons showing co-expression. (C) Representative unit recordings from LC neurons. Insets show action potential waveforms, while raster plots and PSTHs show typical biphasic response to contralateral (left) and ipsilateral (right) toe pinch. (D) Top, representative high-pass filtered (>300 Hz) traces of two simultaneously recorded channels in response to 1 Hz on/off laser stimulation. Middle and bottom, raster plot and PSTH of neuron from bottom trace. (E) Same as (D) for stimulation at 5 Hz (pulse duration = 10ms). (F) Same as (E) for laser stimulation at 10 Hz (pulse duration = 10ms). (G) Laser-evoked firing rate as a function of light intensity (left, 1sec stimulation), pulse duration (middle, at 5 Hz) or stimulation frequency (right, pulse duration 10ms). (H) Representative EEG and EMG traces showing an immediate awakening from NREM sleep upon strong optogenetic LC activation (10 s stimulation at 10 Hz, 90ms pulses). (I) Representative EEG spectrograms after LC activation (left) vs. sham stimulation (right).

In freely behaving rats, strong (10sec at 10Hz, 90ms pulse duration) LC activation reliably led to awakenings from NREM sleep and to EEG activation, in line with previous studies ^21, 22^ (Fig. 3h,i). The ability to elicit short arousals upon unilateral LC optogenetic activation was maintained for 6 months, attesting to sustained and stable CAV-2-mediated ChR2 expression with minimal toxicity (as previously noted ^22, 44^), also validated by immunohistochemistry (Methods).

Next, we asked whether LC opto-activation alters pupil size. Under light anesthesia, LC opto-activation (3 s stimulation, at 10 Hz) induced a large increase in pupil size (32.38%, 74.56%, 107.42%, 145.89% and 198.64% increase compared to baseline for 10, 30, 50, 70 and 90ms illumination pulse durations, respectively; Fig. 4 and Supp. Video 1). Together, these data demonstrate that optogenetic LC excitation elicits reliable behavioral, electrophysiological, and pupillary markers of arousal.

**Figure 4.**
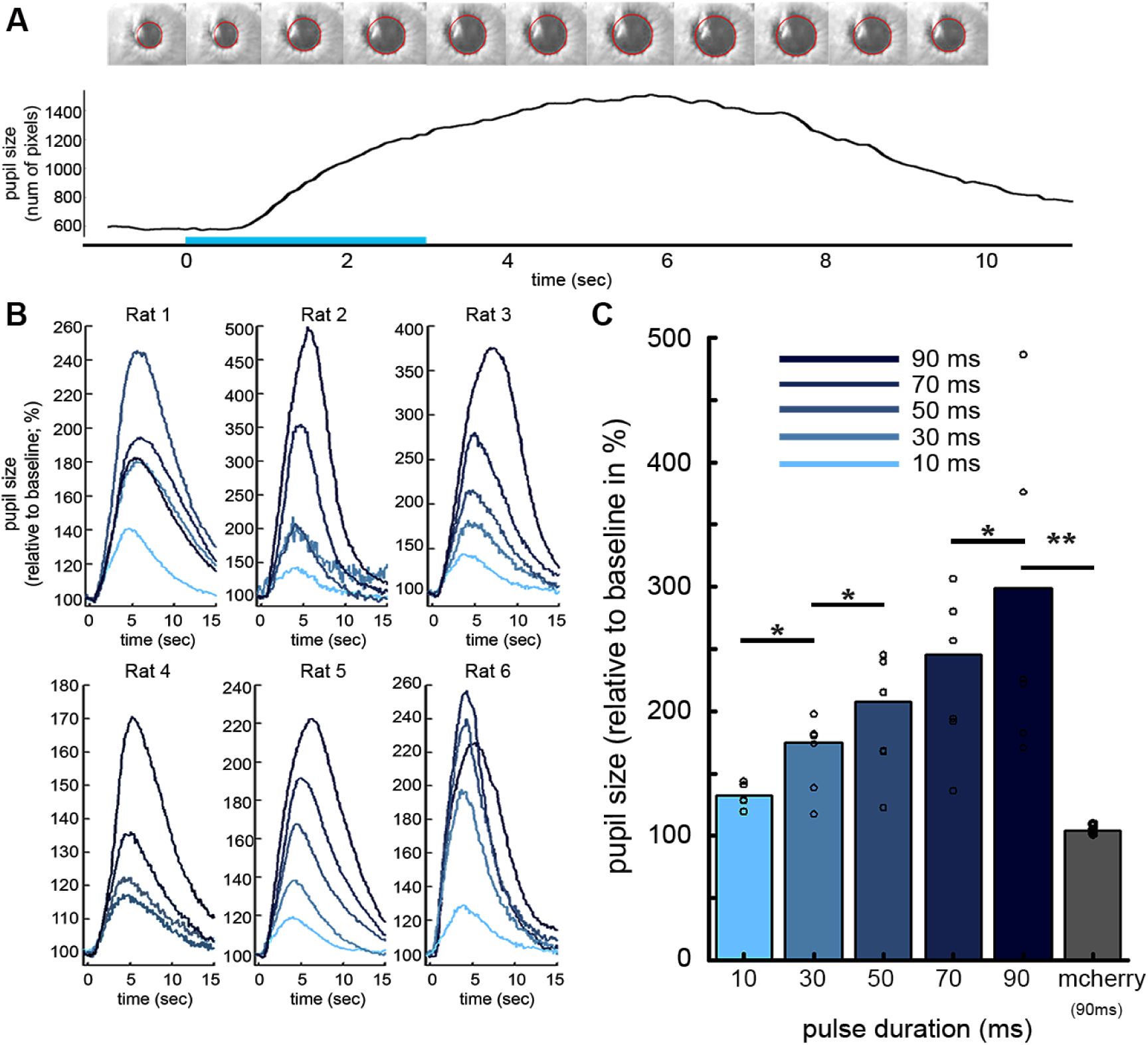
Optogenetic activation of LC neurons causes pupil dilation. (A) Representative single trial example of LC optogenetic induced pupil dilation. Top, video images and pupil size estimation (red contour) before and during 3 s laser illumination (10 pulses/sec, 90ms pulse duration). (B) 15s time-courses of pupil size increase (relative to baseline) for five pulse duration conditions (10-90ms, light to dark blue) in each animal. (C) Summary data (mean ± SEM, n = 6 rats) of pupil size increase as a function of laser pulse duration, (**P<0.01, *p < 0.05 Wilcoxon test) in n=6 rats (ChR-mCherry, in blue) and n=5 control rats (mCherry, in gray).

### Liminal optogenetic LC excitation increases probability of SEAs during NREM and REM sleep

To examine the causal effects of LC activity on arousal threshold during sleep, we combined LC optogenetic activation with auditory stimulation. Pupil dilation (under light anesthesia) and awakenings during NREM sleep from adequate opto-activation were prerequisites for inclusion of rats in the *in vivo* auditory arousal threshold experiments. For each animal, we titrated laser stimulation to identify a liminal level that was insufficient on its own to elicit reliable awakenings (by shortening stimulation duration to 3s and shortening individual pulse duration to 10 – 30ms, see Methods). Next, during the arousal threshold experiment, tone pips were delivered at 67 and 80 dB SPL. In half of auditory trials, sounds were presented on a background of liminal laser illumination. In addition, laser was activated in the absence of sound to evaluate the effect of the laser alone (Fig. 5a). We then compared awakening probability in sound-only trials (“S”), laser-only trials (“L”), and their co-occurrence (“SL”) (Fig. 5a). LC activation significantly increased awakening probability (29.8 ± 5.9% and 25.3 ± 8.3% increase from NREM sleep and from REM sleep, n = 8 rats and n = 6 rats, respectively; Fig. 5b,c and Supplementary Fig. 2). This increase was significantly greater than that expected assuming independent effects of LC activation and auditory stimulation (yellow bars in Fig. 5, Methods). We found a change in SEAs probability for all stimulation regimes (5 Hz, 10 Hz or “phasic mode”, Methods; Supp. Fig. 2). Since sleep spindles are associated with higher arousal threshold during NREM sleep ^45, 46^, we analyzed their occurrence during liminal LC excitation associated with increased SEA probability. Indeed, LC optogenetic activation (at 10Hz) before sounds presented during NREM sleep decreased spindle occurrence (0.052 spindles/sec vs. 0.036 spindles/sec for ‘laser off’ and ‘laser on’ trials, respectively, n=8 rats, p=0.017 via one-tail paired t-test), in agreement with previous studies ^12, 38^

**Figure 5.**
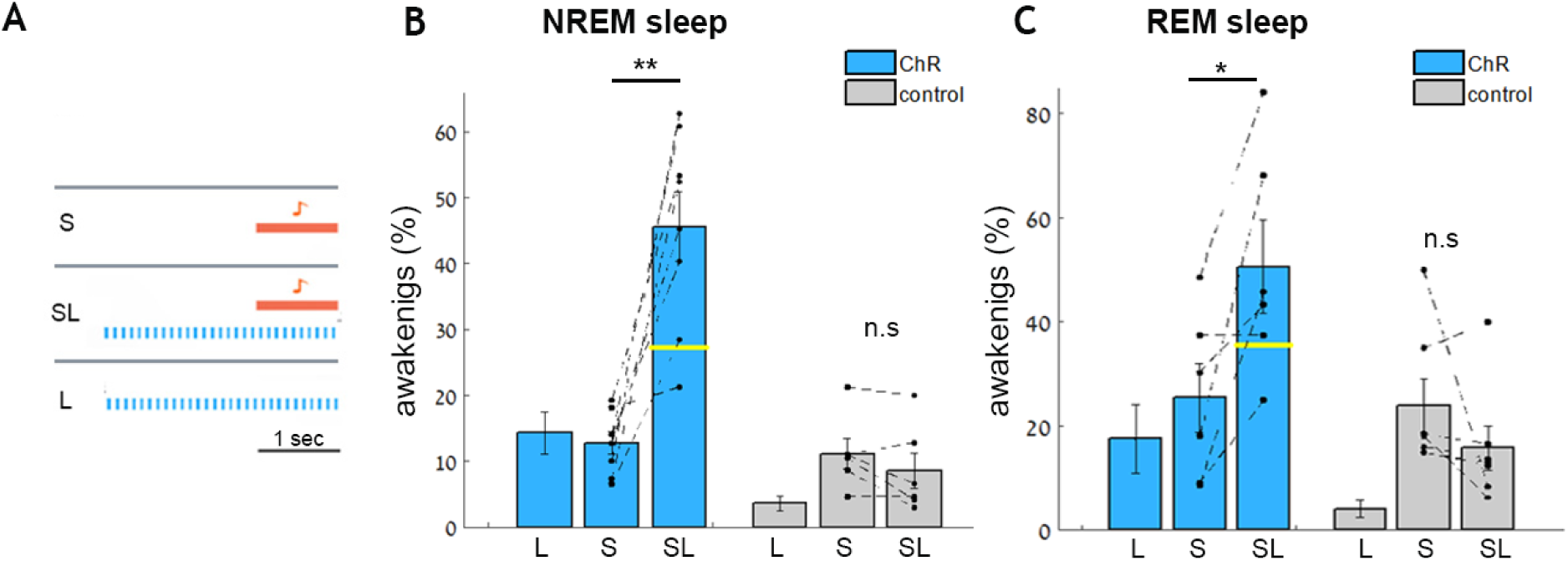
Liminal optogenetic LC excitation increases the probability of sound-evoked awakenings. (A) Schematic of three conditions in LC optogenetic-activation arousal threshold experiments: presentation of sound (S) only, sound on a background of blue laser stimulation (SL), blue laser only (L). (B) Probability of awakening as a function of the three experimental conditions in rats injected with CAV-PRS-ChR2-mCherry (blue, n = 8) or control mCherry (gray, n=6) in NREM sleep. Yellow horizontal line dividing the SL bar represent the expected independent effect of sound and laser stimulation (Methods). (C) Same as (B) for REM sleep with rats injected with CAV-PRS-ChR2-mCherry (blue, n = 6) or mCherry (control, gray, n=6). **p <0.01, *p <0.05, paired t-tests corrected with FDR (see Supplementary Figure 2 for laser frequency parameters).

### Optogenetic silencing with an anion-conducting channelrhodopsin effectively suppresses LC neuronal activity and causes pupil constriction

Multiple wake-promoting systems can lead to sleep-wake transitions, and as we have seen, selective LC-NE activity is sufficient to increase probability of SEA. But does selective silencing of this system alone during sleep decrease SEAs? In other words, is LC-NE activity necessary to modulate arousal threshold? To address this, we specifically and efficiently silenced the LC with high temporal resolution using a CAV-2 vector harboring a soma-targeted anion-conducting opsin stGtACR2 ^47^ under the control of the PRS promoter (CAV2-PRS-stGtACR2-fRed). After bilateral LC injections, we observed specific transgene expression in the LC (Fig. 6a).

**Figure 6.**
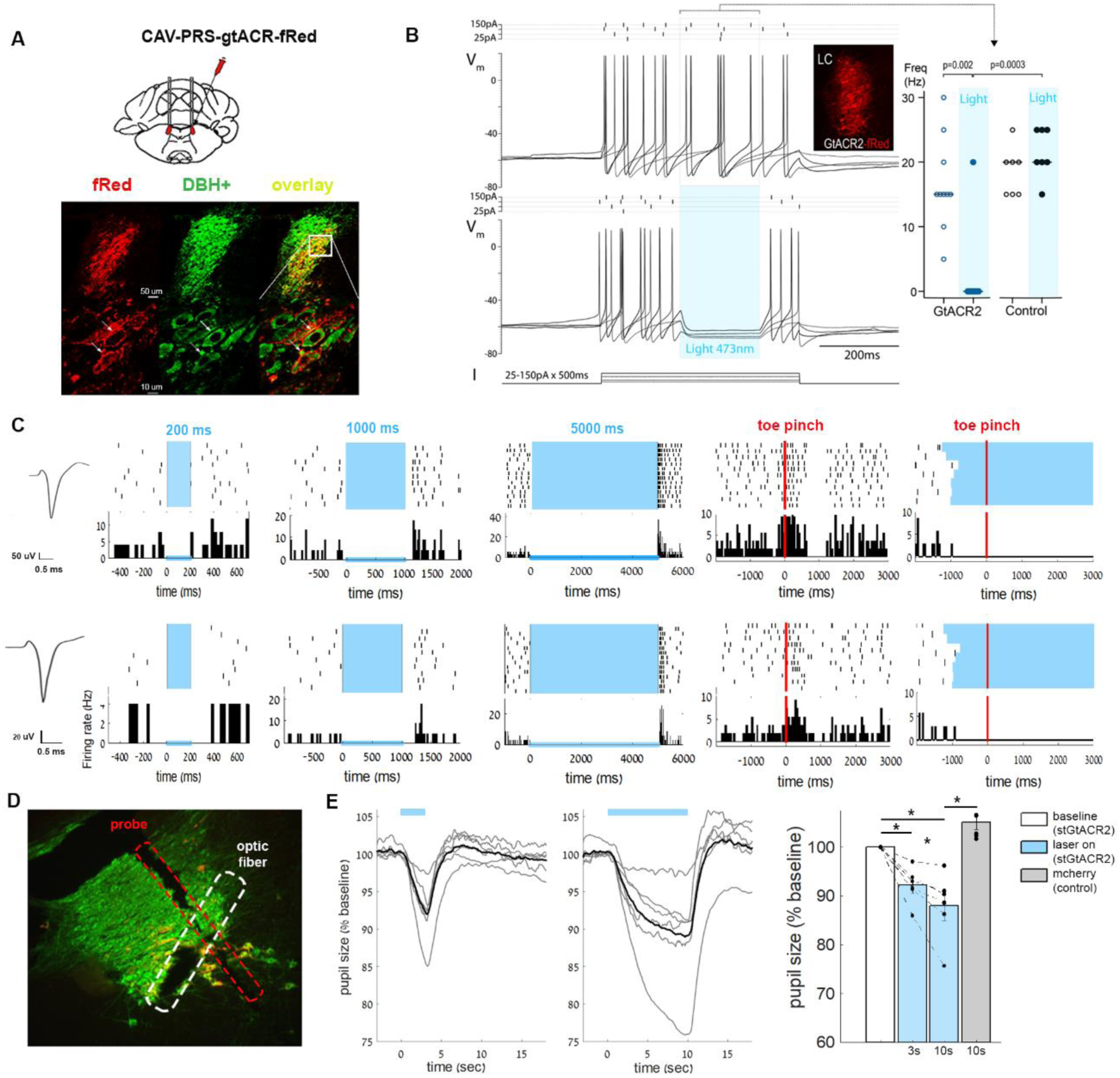
Specific and effective optogenetic inhibition of LC neurons. (A) Schematic of bilateral LC injection and optic fiber implantation (top) and specific expression of stGtACR2-fRed in LC neurons (bottom): representative coronal images showing CAV2-PRS-stGtACR2-fRed expression (left column, red), labeling of TH+ neurons (middle column, green), and their overlay (right column, yellow). (B) Whole-cell current-clamp recording from LC neuron in pontine slice showing the spike discharge evoked by current injection is prevented by a brief pulse of light (473nm, 200ms x 4mW) to activate the stGtACR2, a silencing effect seen across recordings from transduced LC neurons (n=10, data from response to 175pA pulse, p=0.002 Wilcoxon matched pairs) whereas no effect of light was seen in non-transduced LC neurons in the same slices (n=7). Native fRed fluorescence (shown in inset) was easily observed in live slices allowing cell identification. (C) Representative data from two single-unit recordings of neurons in a CAV2-PRS-stGtACR2-fRed transduced rat. (left) action potential waveforms and (middle) raster plots and PSTH of inhibitory responses to different laser durations (in blue, 200ms, 1000ms and 5000ms) and (right) LC neuron with typical biphasic response to contralateral toe pinch, abolished by concurrent illumination. (D) Histology of probe track and optic fiber in LC (DBH+ neurons in green, co-localization of DBH neurons and fRed (vector) in yellow (E) Representative pupil size traces (grey lines represent single trials, black line represents average of the trials) during 3sec and 10sec of laser illumination. Bar graph shows significant pupil constriction when LC was silenced in a duration-dependent manner (stGtACR2 in blue, n=6; control mcherry in gray, n=6), Wilcoxon test, *p <0.05.

To evaluate the functional effects of stGtACR2on LC neurons *in vitro* we transduced rats (p21) unilaterally with CAV2-PRS-stGtACR2-fRed, and 2-3 weeks later prepared acute pontine slices for whole-cell patch clamp recordings. Recordings from transduced LC neurons (n=14) showed that opto-activation of stGtACR2 silenced spontaneous firing by activating a potent shunting conductance that reversed close to the calculated chloride reversal potential (Supplemental Fig. 5). This was capable of blocking action potential discharge evoked by intracellular current injection (driving firing to ∼15-20Hz) with rapid onset and offset (Figure 6b).

To stringently test the effectiveness of the stGtACR2 silencing strategy *in vivo* we transduced adult rats unilaterally with a relatively low titer of CAV2-PRS-stGtACR2-fRed (1.4×10^7^pp) and 3-4 weeks later recorded LC action potential discharge under urethane anesthesia with 32-channel silicon probe. Laser illumination of the LC, delivered via a separate optic fiber, (445nM, 10mW at end of fiber) strongly silenced neurons, including during trials when paw pinches or foot-shocks were delivered (Fig. 6c).

Such LC opto-silencing led to robust pupil constriction even during light anesthesia, when pupils are already constricted at baseline. Continuous bilateral laser stimulation significantly decreased pupil size compared to baseline (8.2 ± 3.66% and 14.9 ± 6.9 % decrease for 3s and 10s laser pulses, respectively; Fig. 6e, n=6). Pupil constriction was parametrically dependent on laser stimulus duration whereby longer illumination (10s vs. 3s) led to a significantly (p<0.05) larger effect. The average time to nadir was 3.14 ± 0.28s after laser onset for 3s stimulation, and 9.26 ± 0.79s for 10s stimulation. Thus, CAV-2-mediated expression of stGtACR2 allows effective opto-silencing of LC neuronal activity which causes pupil constriction.

### Optogenetic LC silencing decreases probability of SEA from NREM sleep

To assess whether LC silencing affects arousal thresholds during sleep, we combined LC optogenetic silencing via stGtACR2 with auditory stimulation. To ensure effective LC silencing in each animal, pupil constriction in both eyes (under anesthesia as above) was a prerequisite for *in vivo* auditory arousal threshold experiments. Laser stimulation was applied for 5s continuously (5-7mW at the end of the fiber), either simultaneously with the sound (SL) or, motivated by the difference in baseline tonic effects noted previously (Fig. 2) starting 2s before sound onset (SafterL) (Fig. 7a). LC silencing significantly increased EEG SWA in NREM sleep (Supplementary Fig. 3h) as was reported for LC silencing during wakefulness ^21^.

**Figure 7.**
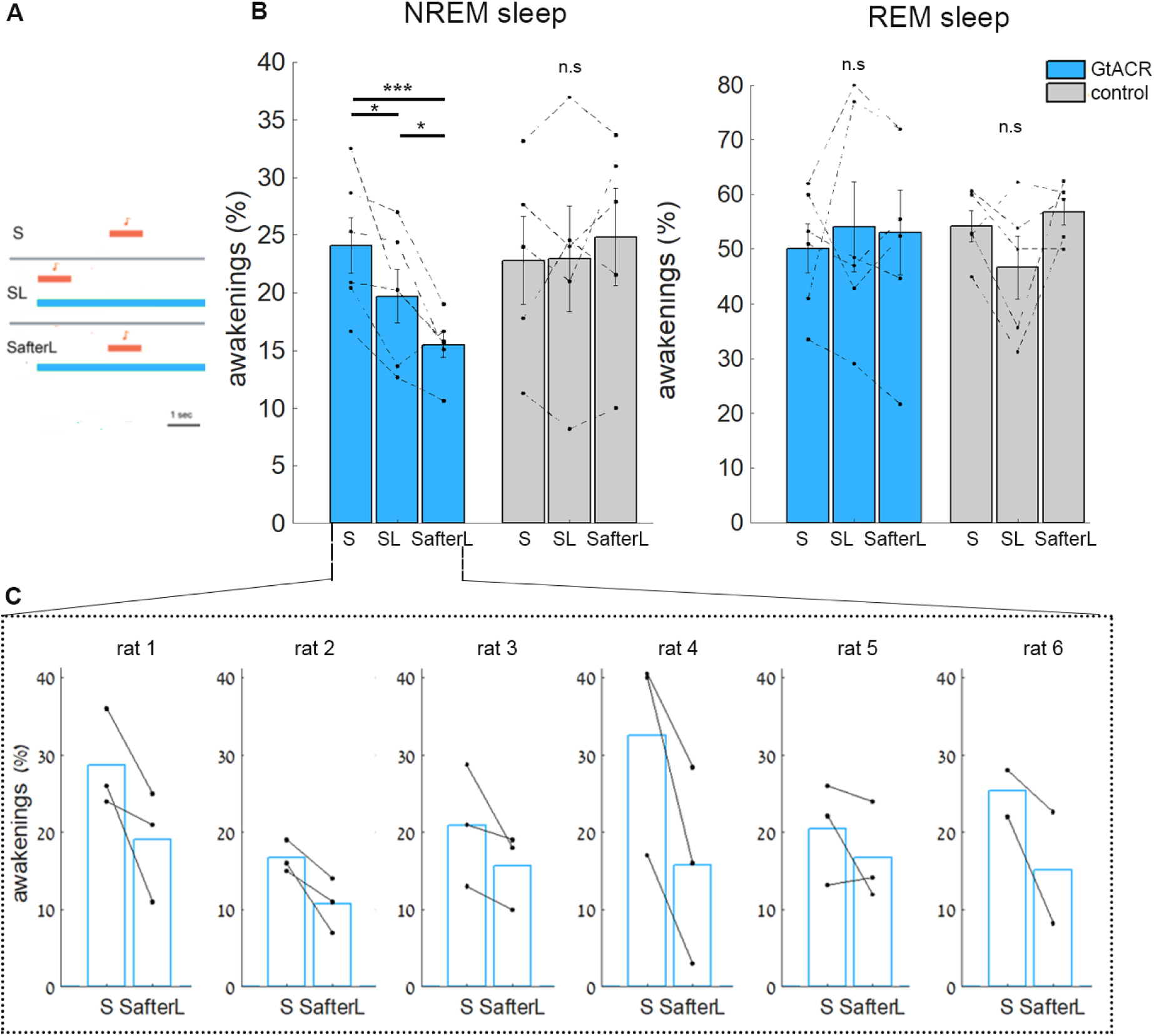
Optogenetic LC silencing decreases the probability of sound-evoked awakenings. (A) Schematic of three conditions in LC opto-silencing arousal threshold experiments. Top, presentation of sound (S) only; middle, sound played simultaneously with laser stimulation (SL); bottom, sound played two seconds after laser onset (SafterL). Laser duration was continuous for 5s. (B) Probability of awakening as a function of the three conditions in rats expressing stGtACR2-fRed (blue, n=6) and control (grey, n=5) in NREM sleep (left) and in REM sleep (right). (paired t-test*p <0.05, **p <0.01). (C) Probability of awakenings from NREM sleep in each session per rat expressing stGtACR2-fRed.

SEA probability during NREM sleep (but not REM sleep) significantly decreases in both conditions (4.38 ± 2.01% and 8.59 ± 2.10% decrease for SL and SafterL, respectively, n = 6) with a significantly more pronounced decrease when baseline LC activity was silenced before sound onset (Fig. 7b,c). Unlike the association between LC excitation and reduced spindles, LC silencing did not significantly affect spindle occurrence, although there was a trend towards increased spindle events (p=0.068 via one-tail paired t-test, n=6).

## Discussion

After validating an experimental paradigm to quantify auditory arousal threshold in freely sleeping rats, we tested the link between LC-NE activity and SEAs. We established a relationship between baseline tonic LC activity and the probability of SEA, and found that pharmacologically decreasing NE signaling reduced SEA probability. Liminal optogenetic LC excitation increased SEA probability during both NREM and REM sleep, while optogenetic silencing decreased SEA probability during NREM sleep, especially when silencing began shortly before sound onset. These results causally link activity in the LC to sensory stimulus-evoked awakening and support the idea that the absence of LC activity during sleep is responsible for the elevation of arousal thresholds.

### Mechanisms controlling sensory-evoked awakening

Multiple overlapping neuromodulatory systems are implicated in sleep-wake regulation, including noradrenaline, histamine, acetylcholine, hypocretin, and dopamine ^26,48,49^. Strong activation of any of these systems promotes arousal whereas their inhibition reduces wakefulness. Our results go beyond intrinsic mechanisms controlling awakening and highlight a key role for the LC-NE system in mediating sensory-evoked awakening. Given the overlap in neuromodulatory activities, other mechanisms likely also contribute to this process, as was shown for dorsal raphe dopamine activity ^23^. Wakefulness does not depend on one exclusive system, because even simultaneous lesions of histaminergic, noradrenergic, and cholinergic neurons do not robustly affect daily levels of wakefulness ^50^. Given that being able to wake up in response to external stimuli is crucial for survival, overlapping mechanisms likely contribute.

The mechanisms downstream from LC that modulates SEAs remain unclear. Our optogenetic experiments targeted LC cell bodies, and future studies are needed to determine the roles of NE (and dopamine) release at specific projection targets. While LC has been typically regarded as a relatively homogeneous nucleus ^51^ some recent studies challenge this view and suggest that LC subpopulations have distinct projections that serve different functions ^30,40,41,52–54^. Downstream effects of LC on SEAs could be mediated through direct cortical projections, where α-adrenergic receptors mediate cortical desynchronization ^55, 56^. The LC may also act through thalamic projections ^56^. Infusion of NE (or LC stimulation) increases thalamic excitability ^57^, suppresses ongoing rhythmic activity ^58, 59^, and accelerates emergence from anesthesia ^60, 61^. Another possibility is that LC affects other modulatory centers such as the basal forebrain ^62–64^, where it activates cholinergic wake-promoting neurons ^65^ and inhibits GABAergic sleep-promoting neurons ^66^. Indeed, injection of NE in the basal forebrain of sleeping rats causes SWA suppression and promotes awakening ^67–69^. LC projections to the hypothalamus, the amygdala or other regions could also be involved.

### A causal link between LC activity and pupil size

Pupil size is a marker of arousal ^70, 71^ and is often used to indirectly infer LC-NE activity levels across multiple species ^72–75^. Pupil size is tightly correlated with LC activity in monkeys ^76, 77^ and humans ^78^ and with cortical noradrenergic activity in mice ^79, 80^. In addition, electrical stimulation of LC causally affects pupil size ^76, 81^. Here, cell-type-specific LC optogenetics establishes a robust causal bidirectional influence of LC activity on pupil size. Under light anesthesia, LC optogenetic activation drives large pupil dilation, while optogenetic silencing causes a modest but consistent pupil constriction, possibly reflecting a ‘floor’ effect related to anesthesia. Our results in rats support a recently published study in mice ^80^ in showing that LC activity is sufficient to bidirectionally alter pupil dilation. LC-mediated pupil dilation may act through disinhibition of parasympathetic pathways (e.g. via cholinergic pre-ganglionic neurons of the Edinger-Westphal nucleus controlling the sphincter muscle) ^77, 82^ with possible contribution of sympathetic pathways controlling the iris dilator muscle ^81,83,84^. Our results provide a robust experimental framework to explore the precise mediating pathways in detail.

### Tonic and phasic LC activity

The LC shows two types of activity: a tonic mode that is high in wakefulness and low in sleep ^12^ implicated also in arousal, stress, anxiety, and pain processing ^85^; and a phasic mode that occurs during presentation of salient stimuli ^29,86,87^, behavioral tasks ^88^ and reward anticipation ^89^. Our results highlight a role for tonic (but not phasic) LC activity in SEAs. For example, SEA probability was significantly lower when LC silencing started before sound presentation than when LC silencing began simultaneously with the sound (Fig. 7b). In contrast, phasic sound-evoked LC responses were not linked to SEAs and were inversely proportional to levels of baseline tonic LC firing. The link between tonic LC activity and SEAs observed here, together with the link between tonic LC activity and anxiety ^31, 90^, suggests how anxiety may promote awakenings ^91, 92^. An active inference model recently proposed that LC activity represented a state-action prediction error where uncertainty was reflected in increased LC activity which in turn promoted behavioural flexibility in behaving agents ^93^. This may be extended to sleep states where the observed tonic LC activity represents uncertainty about internal state and influences the probability of response (awakening) to a sensory stimulus (sound). Along this line, LC-NE activity may be an important factor driving variability in arousal threshold across individuals. We were surprised that phasic LC activity was not related to SEA probability, because it is reported to be related to salient stimuli. Thus, the mechanisms that underlie more frequent awakening in response to behaviorally relevant stimuli ^6, 7^ remain to be elucidated. While the current results indicate that LC activity robustly modulates SEA, it remains unclear to what extent this co-occurs with corresponding response modulations along sensory pathways. While LC-NE signaling clearly modifies response fidelity across sensory modalities and species ^49,94–96^, it is unlikely that it simply “gates” propagation of auditory signals to the cortex as auditory responses in primary cortex are largely comparable across vigilance states ^97–99^. Instead, tonic LC activity that drives SEAs may facilitate sensory processing downstream from primary regions, by acting at higher-order regions or non-sensory pathways to promote awakenings.

In summary, we show that low LC activity during sleep plays a key role in mediating reduced responsiveness to sensory stimuli, thereby revealing a mechanism that controls how sleep is maintained in the face of external events.

## Acknowledgements

We thank Inna Slutsky and members of Nir lab for discussions and suggestions, Chiara Cirelli and Giulio Tononi for comments on earlier draft, Yaniv Sela for help with spindle detection, Menahem Segal and Susan Sara for helpful discussions on LC experiments. We thank the Sackler Cellular & Molecular Imaging Center (SCMIC) and Dr. Leonid Mittelman for dedicated assistance. This work was supported by the Israel Science Foundation (ISF) grants 1326/15 and 51/11 (I-CORE cognitive sciences) and the Adelis Foundation (YN). EJK is an INSERM fellow. OY is supported by the European Research Council (ERC-2013-StG OptoNeuromod 337637) and the Adelis Foundation. CAV-2 vector production was supported by CNRS BioCampus (Montpellier). AS is a Wellcome Trust funded PhD student on the Neural Dynamics program.

## Author contributions

H.H., A.E.P. and Y.N. conceived research and designed experiments.

H.H., N.R., and N.M. performed surgeries, collected and analyzed data, with supervision from Y.N.

A.S. and E.P.R. performed surgeries, collected and analyzed in-vitro and acute LC silencing data, with supervision from A.E.P.

A.J.K. and L.B. assisted with surgeries and auditory stimulation aspects.

M.L., Y.L., A.E.P. and E.J.K. generated CAV2-PRS-Cre-v5 and CAV2-PRS-GtACT2-fRed.

O.Y. and A.E.P. provided optogenetic constructs and expertise.

H.H. and Y.N. wrote the manuscript.

All authors provided ongoing critical review of results and commented on the manuscript.

## Online Methods

### Animals

All experimental procedures at the University of Tel Aviv including animal handling, surgery, and experiments followed the NIH Guide for the Care and Use of Laboratory Animals, and were approved by the Institutional Animal Care and Use Committee (IACUC). All experimental procedures at the University of Bristol conformed to the UK Animals (Scientific Procedures) Act 1986 and were approved by the University Animal Welfare and Ethical Review Board. Adult male Long-Evans rats (Supplier e.g. RRRC, 8–12 weeks old at the time of surgery) were used for arousal threshold, pharmacological, and optogenetic experiments, except for acute opto-silencing where adult Lister Hooded rats were used (supplied by Charles River, 9-12 weeks old at time of surgery). Adult male Wistar rats (Supplier Harlan Israel, 8–12 weeks old at the time of surgery) were used for chronic recordings. Rats were housed individually in transparent Plexiglas cages at constant temperature (20°-23°C), humidity (40-70%) and circadian cycle (12h light/dark cycle, starting at 10:00 AM). Food and water were available ad libitum. All behavioral and electrophysiological experiments were performed between 10 a.m. and 10 p.m. A few days prior to surgery, rats were placed in their home cage for habituation to the experimental environment.

### Viral Vectors

Viral vectors included CAV2-PRS-hChR2(H134R)-mCherry for LC opto-excitation ^22^, and CAV2-PRS-stGtACR2-fRed (for LC opto-silencing – based on ^47^). CAV-2 vector titers were between 5 to 14 × 10^12^ physical particles/ml and 1 x 10^9^ PP were typically injected to the LC. For control animals a double injection of CAV2-PRS-Cre-V5 (x10 dilution) and AAV5-EF1a-DIO-mCherry (Viral Core Facility, University of Zurich), in a 1:1 ratio. The generation and validation of CAV2-PRS-ChR2-mCherry for selective LC opto-excitation is described in our previous work ^22^. In brief, a 240-bp PRSx8 synthetic promoter sequence was used ^100^ that restricts the expression of the transgene to a subset of neurons that express the Phox2 transcription factor, which are predominantly catecholaminergic ^101, 102^. CAV2-PRS-stGtACR2-fRed was generated by the Platform de vectorology de Montpellier (PVM) using methods described in del Rio et al (submitted). Details of the construction are available on request: briefly, CAV2-PRS-stGtACR2-fRed contains the PRS promoter driving stGtACR2 in sequence with fusionRed (fRed), a monomeric fluorescent protein ^103^, and a poly-adenylation signal (polyA) from SV40. This cassette replaces the E1 region in a E1/E3-deleted, replication-defective CAV vector. Similarly, CAV2-PRS-Cre-V5 was generated in the Pickering Lab and has the same promoter with Cre-recombinase and a V5 tag in the expression cassette.

### Surgery

Before all surgical procedures, rats were anesthetized (with either isoflurane 4% by volume for induction and 1.5% for maintenance or with ketamine (50 µg/kg, Vetalar; Pharmacia, Sweden) and medetomidine (300 mg/kg, Domitor; Pfizer, UK)) and placed in a stereotaxic frame. Surgery was performed in sterile conditions, following procedures described in ^53, 98^. Under microscopic control, a craniotomy was made using a high-speed surgical drill. When a probe/optrode was implanted, the dura was carefully opened. Then electrodes or injection needles were advanced into the brainstem after penetrating the pia mater while carefully avoiding vasculature. Two-component silicone gel (KwikSil; World Precision Instruments, FL, USA) was applied to seal the craniotomy and protect the brain surface. After gel polymerization, dental acrylic was gently placed around the electrodes or optic fibers, fixing them to the skull. EEG screws were placed over the frontal and parietal cortices. Ground and reference screw electrodes were placed above posterior parietal lobe (ChR2 and associated experiments as well as chronic LC recordings) or above cerebellum (otherwise). Neck muscle electrodes were implanted bilaterally and bipolar referencing used for electromyography (EMG). A custom-made EEG/EMG connector was fixed on the skull with dental cement for sleep recordings.

For all experiments (except for acute silencing recordings), viral vectors were injected to the LC (using an UMP3 Microsyringe Injector and Micro4 Controller pump, and 33G Nanofil needles WPI, USA) unilaterally (for activation) or bilaterally (for silencing) at coordinates: anteroposterior (AP): λ −3.7 mm; medio-lateral (ML) +/−1.18 mm, dorso-ventral (DV) 6.2-6.6 mm with rostral angulation of 15°, and at a rate of 100 nL/min for 12 min (3 injections of 400 nL for a total volume of 1.2 µl). For light stimulation/inhibition an optic fiber (200-µm diameter 0.22 NA) was implanted (MFC_200/240-0.22_10 mm_MF1.25_FLT, Doric Lenses) above the LC. Unilateral and bilateral fiber implants were used for LC opto-excitation and opto-silencing respectively, as was the case for virus injections, at these coordinates (AP: λ −3.7 mm; ML +1.21 mm; DV 5.4 mm with rostral angulation of 15°). In acute silencing experiments, viral vector was injected to the LC (using a Neurostar injector-drill robot on a Kopf stereotaxic frame) at coordinates: AP: λ −2.1 mm, ML: +1.2 mm, DV: 5.3-5.8 mm with rostral angulation of 10°.

### Electrophysiology and LC recordings

#### Brain slice recordings

Male Wistar rats were used for slice electrophysiology using previously described methods (Hickey *et al.* 2014 & Hirschberg *et al.* 2017). CAV2-PRS-stGtACR2-fRed was injected (0.69-1.38×10^9^ PP in total) under recovery anesthesia (ketamine (50 mg/kg, Vetalar; Pharmacia) and medetomidine (300 μg/kg, Domitor; Pfizer)) at postnatal day 21. After burr hole formation a dorso-ventral injection track for a vector-filled, pulled glass pipette (tip diameter 20-30µm) was started at 1mm lateral and 1mm posterior to lambda with 10° rostral angulation. Four injections of 250nl of vector were made at depths from −4.6 to −5.5mm unilaterally to the LC. Rats were terminally anesthetized with isoflurane (5%) 14-21 days later, decapitated and the brain quickly removed and immediately chilled in ice-cold cutting solution (similar to the recording solution but NaCl was reduced to 85 mM and substituted with sucrose 58.4mM). Transverse slices (300–350μm thick) of the pons were cut from dorsal to ventral using a vibratome (Linearslicer Pro 7; DSK) in cold (4°C) cutting solution.

After cutting, slices were kept at room temperature in carbogenated recording solution (NaCl 126, KCl 2.5, NaHCO_3_ 26, NaH_2_PO_4_ 1.25, MgCl_2_ 2, CaCl_2_ 2, and D-glucose 10 (in mM) saturated with 95%O_2_/5%CO_2_, pH7.3, osmolality 290mOsm/L) for at least 1 h to recover before recording. Pontine slices were transferred into the recording chamber of an upright fluorescence microscope (DMLFSA; Leica Microsystems), superfused with artificial CSF at a rate of 4–8 ml/min heated to 35°C. Borosilicate glass patch pipettes (Harvard Apparatus GC120F-10, resistances of 3–6MΩ) were filled with internal solution (K-gluconate 130, KCl 10, Na-HEPES 10, MgATP 4, EGTA 0.2, and Na_2_GTP 0.3, (in mM)). Cells were identified under gradient contrast illumination and examined for transduction by epifluorescence illumination. Blue light (473nm, Doric LED, 4mW) was pulsed onto slices from an optical fiber (400um diameter) placed above the LC. Whole cell voltage and current clamp recordings were obtained with a Multiclamp 700A amplifier (Axon instruments). All membrane potentials were corrected for a junction potential of 13 mV. The reversal potential for chloride ions using this combination of internal and external solutions was calculated as −68.7mV. Data was acquired using a Power1401 A-D converter (CED) and displayed and analyzed with Spike2 software (CED, Cambridge Electronic Design).

#### Chronic experiments

In all chronic experiments, epidural EEGs (filtered 0.1-100 Hz), muscle tone EMG (filtered 10-100Hz) and infrared video (except for in optogenetic experiment) were continuously recorded using a TDT RZ2 electrophysiology system, PZ amplifier and RV2 video capture system (for pharmacological experiments) from Tucker Davis Technologies, USA. For chronic LC recordings, a 16-channel linear silicon probe (177 µM^2^ contact surface area, 100 µM between adjacent contacts, Neuronexus, USA) mounted on a micro-drive was implanted above the LC (AP: λ −3.7 mm; ML +1.15-1.18 mm). After recovery and auditory habituation, the chronic micro-drive electrode was lowered 50-200μm daily under light isoflurane sedation until we succeeded in recording from neurons responding to toe-pinch stimulation.

#### Acute recordings

For acute opto-excitation recording, three weeks after viral vector injection, an optrode was lowered progressively in the LC (same electrode specifications as in chronic experiment (see above), coupled with a 100µm core diameter optic fiber mounted 100µm above the top electrode contact; Neuronexus, USA).

For acute opto-silencing recordings, rats were anesthetized with urethane at a dose of 1.5mg/kg body weight and fixed in a stereotaxic frame. Isoflurane (at 1-2%) was also initially used as a supplement to urethane for the craniotomy surgery. Two craniotomies were made, each consisting of a ‘window’ of side length between 0.5-1mm centered on target coordinates from lambda AP:-2.1 mm, ML: 1.2 mm and AP: +1.4, ML: 1.2 mm. A tapered fiber optic (Optogenix, type Lambda-B) was inserted via the caudal window at a 10° rostral angulation to a depth of ∼5.5 mm from the surface of the brain. A 32-channel silicon recording probe (Neuronexus, A1x32-Poly2-10 mm-50 s-177) was then inserted using a Narishige hydraulic manipulator (MWS31) via the rostral craniotomy, at an angle of −20° to depths of up to 8mm from brain surface. The probe signal was digitized using an 32-channel headstage (RHD2132, INTAN); the resulting signal was acquired and displayed using the OpenEphys recording system ^104^. LC neurons were identified initially by their pattern of spontaneous discharge (0.2-5 Hz), large amplitude long duration action potentials and their biphasic response to hindpaw pinch. Once the LC was identified, then the field was illuminated (445nm, 15mW at the tip, OMICRON PHOXX laser) and the effect on spike discharge examined. Post hoc histological examination confirmed the position of the recording probe in the LC.

### Optogenetic laser parameters

Blue stimulation at 447 nm for optogenetic excitation and silencing was delivered via lasers (CNI, China) coupled to optic fibers whose timing and intensity were automatically controlled via RZ2 (TDT). Light intensity at optic fiber tips was measured with a power meter (ThorLabs PM100D) prior to optic fiber insertion. Before starting opto-activation arousal threshold experiments (Fig. 5), we confirmed effective LC opto-excitation in each animal by verifying that strong laser stimulation (10 s at 10 Hz, 90ms pulse duration, 20 mW) evoked reliable EEG desynchronization and sleep-wake transitions (Fig. 3h,i), as shown in ^21, 22^. Only rats that woke up from sleep in response to strong laser stimulation were included in subsequent experiments with auditory stimulation. Next, we adjusted the laser stimulation parameters (total duration 3sec, pulse duration 10-30ms), such that laser stimulation alone did not reliably affect awakening probability.

### Auditory stimulation

All experiments were conducted in a double-wall sound-attenuating acoustic chamber (Industrial Acoustics Company, Winchester, UK). All sounds were stereo signals programmed in Matlab, where one channel (containing a 1s 4 kHz tone pip) was routed to the mono speaker in the chamber and the other channel was routed to the electrophysiology acquisition system. The sounds were transduced into voltage signals by a high-sampling rate (192 kHz) sound card (LynxTWO, Lynx, USA), amplified (SA1, Tucker Davis Technologies (TDT)), and played free-field through a magnetic speaker (MF1, TDT) mounted 35cm above the animal. For optogenetic experiments, where precise synchronization between auditory and laser stimulation was required, sounds were digitally loaded from a PC and routed to the mono speaker via a TDT RZ2 system (Tucker Davis Technologies, USA). Tone pips played during arousal threshold experiments were with intensities in the range of 50-98 dB SPL with intervals of 55s to 105s with ± 15 s jitter (the louder were the sounds, the higher was the interval between sounds). Sound intensity levels (dB SPL) were measured at cage floor height.

### Systemic pharmacology

For systemic pharmacology experiments (Fig. 1d,e & Supp. Fig. 1) either detomidine (1mg/Kg; α2 agonist to reduce NE levels), yohimbine (1 mg/Kg; α2 antagonist to increase NE levels) or saline (0.9% NaCl; 1 ml/g) were injected i.p. in pseudo-random order while rats were lightly sedated with isoflurane. Recordings and acoustic stimulation started 15-20 min after the injection.

### Combined arousal threshold & optogenetic experiments

In opto-activation experiment, we used three separate experimental paradigms (on different days) to evaluate the effects of laser stimulation with pulses applied at either 5 Hz, 10 Hz, or “burst mode” where 10 pulses were delivered at 20 Hz, repeated every second for 3s. Sounds consisted of 1s 4kHz tone pips presented at either 67 dB or 80 dB SPL. Each experiment lasted ∼12h and included three conditions (Fig. 5a): (i) A “sound-only” condition (“S”) where tone pips were presented alone. (ii) A “sound with laser” condition (“SL”) where tone pips were presented during the last second of laser stimulation. (iii) A “laser-only” condition (“L”) where laser stimulation was delivered without sounds.

In opto-silencing experiment, each experiment included four conditions: (i) A “sound-only” condition (“S”) where tone pips were presented alone. (ii) A “sound after laser” condition (“SafterL”) where tone pips were presented two seconds after laser onset (iii) A “sound with laser” condition (“SL”) where tone pips were presented simultaneously with the laser onset (iv) A “laser-only” condition (“L”) where laser stimulation was delivered without sounds. Laser was applied for 5sec continuously. Typically, ∼600 trials (all conditions confounded) occurred in each experiment. Each condition (S, SL, SafterL and L) was compared to the other condition of the same day session.

### Pupillometry

Rats were anesthetized with 1.5% isoflurane and placed in a stereotaxic frame. An optic fiber connected the implanted optic fiber ferrule to the 473nm laser. The eye was illuminated with an infrared light and monitored continuously with a vision color camera (VGAC; TDT) synchronized to a dedicated RV2 video processor (TDT) allowing frame-by-frame synchronization between video data (30 frames/sec) and laser stimulation. We examined (Fig. 4) the effects of different pulse durations (10-90ms) on pupil size. For LC activation, we applied unilateral laser stimulation to the LC while recording from the contralateral pupil. For LC silencing, we applied bilateral laser stimulation while recording from the left eye followed by the right eye and then averaged the pupil size change of the two eyes. In control experiments, we applied unilateral (control for opto-excitation) or bilateral (control for opto-silencing) laser stimulation to the LC.

### Histology

#### Tissue collection and fixation

Following all experiments (except acute silencing experiments), rats were perfused intracardially with saline (0.9% NaCl; 1 ml/g) followed by 4% paraformaldehyde (Merck) under deep isoflurane anesthesia. Brains were then extracted and fixed for 24 - 48 h in 4% PFA. Coronal brain sections were cut using Leica VT1000 S vibrating blade microtome at 50µm. Then, sections were either kept free floating in PBS for immunohistochemistry (IHC) or mounted on glass slides and examined under bright field microscopy. Following acute silencing experiments, rats were killed with an overdose of pentobarbital (Euthatal, 20mg/100g via IP injection) and perfused transcardially with 0.9% NaCl following by 4% formaldehyde in 0.1M phosphate buffer (PB). Brains were removed and postfixed overnight before cryo-protection in 30% sucrose in 0.1M PB. Sagittal tissue sections were then cut at 40µm intervals using a freezing microtome.

#### Immunohistochemistry

The viral vector expression efficacy and specificity was evaluated histologically by double staining of free-floating sections. To this end, following all experiments (except acute silencing experiments), sections were washed 3 times in PBS (phosphate buffered saline, Hylabs) and then permeabilized in PBST (phosphate buffered saline containing 0.1% Triton X-100 (Merck)). Next, sections were blocked in PBST containing 20% NGS (normal goat serum (Vector Laboratories)) for 1 h at room temperature and incubated with primary antibodies in PBST (containing 2% NGS) at 4°C for 24-36 h. Primary antibodies were either against tyrosine hydroxylase (chicken anti-TH 1:500; ab76442 abcam) and mCherry (mouse anti-mCherry, 1:500; ab125096 Abcam) for activation or against dopamine beta hydroxylase (mouse anti-DBH 1:500; MAB308 Merck Millipore) and fRed (rabbit anti-tRFP 1:500; AB233 evrogen) for silencing. After 3 washes in PBS, sections were incubated with secondary antibodies conjugated to fluorophores (either Alexa Fluor488 goat anti-chicken 1:1000; ab150173 abcam and Alexa Fluor594 goat anti-mouse 1:1000; ab150120 abcam for activation or Alexa Fluor488 goat anti-mouse 1:1000; ab150117 abcam and Alexa Fluor594 goat anti-rabbit 1:1000; ab150080 abcam for silencing) in PBST containing 2% NGS for 1.5 h at room temperature. After 3 washes in PBS and once in PBST, sections were mounted onto glass slides and cover-slipped with aqueous mounting medium (Thermo Scientific, catalog # 9990412). Following acute silencing experiments, sections were mounted on slides and permeabilized in 0.1M PB containing 0.3%Triton X-100 (Sigma) then blocked for 45 minutes in 0.1MPB containing 0.3% Triton X-100 (Sigma) and 5% normal goat serum (Sigma). Sections were then incubated on-slide with primary antibodies against dopamine β-hydroxylase (DBH) (mouse anti-DBH, 1:2000, Millipore (Chemicon), MAB308) and fRed (rabbit anti-tRFP 1:500; AB233 evrogen) in PB containing 5% goat serum and 0.3% Triton X-100 for 16 hours. After removal of primary antibodies, tissue sections were washed twice in PB and then incubated with appropriate secondary antibodies conjugated to fluorophores (Alexa Fluor488 goat anti-mouse and Alexa Fluor 594 goat anti-rabbit, both at 1:1000 dilution) in PB containing 0.3% Triton X-100 and 5% NGS at room temperature for 4 hours. Sections were again washed twice in PB and coverslipped with FluorSave mounting medium.

#### Fluorescence Microscopy

Images were acquired by LEICA STED high resolution laser scanning confocal microscope (Leica, Wetzlar, Germany) and a 20 oil/1.4 NA objective. Quantification of extent of co-localization was carried out by Imaris software (Bitplane, Zurich, Switzerland). For histology of rats used in acute silencing experiments, images were acquired by Zeiss Axisoskop2 with pE2 LED excitation system (CoolLED, UK) and analyzed using a Zeiss axiocam and Axiovision software. For tissue from acute optogenetic silencing experiments, images were acquired using a Leica DMI6000 inverted epifluorescence microscope equipped with Leica DFC365FX monochrome digital camera and Leica LAS-X acquisition software.

### Data Analysis

#### Sleep staging and awakenings

Wakefulness was defined by low amplitude, high frequency EEG activity, co-occurring with high tonic EMG activity with occasional phasic EMG bursts. In chronic LC recording experiments, we divided the wakefulness epochs between active wake (grooming, eating, exploration) and quiet wake, based on video recording. NREM sleep was defined by high amplitude, low frequency EEG, co-occurring with reduced EMG tone. REM sleep was defined by low amplitude, high frequency EEG dominated by posterior theta activity co-occurring with flat EMG, as in ^98^.

In arousal threshold experiments, each experimental trial was visually inspected and those occurring during sleep were categorized as either eliciting behavioral awakening or maintained sleep. Behavioral awakening was declared if wake-like EEG flattening (without dominant theta) was present within 3s from sound onset and lasted for at least 3s. A significant percentage of such intervals were accompanied by EMG changes (Supp. Fig. 3), but these did not constitute a necessary condition since changes in neck EMG were not always present, even in cases where video showed clear awakening. When laser was presented alone, behavioral awakening was declared if the changes described above occurred in the same 3s. Arousals could be followed by a wake period or by return to sleep (Fig. 1b). All other trials were categorized as maintained sleep.

#### Arousal threshold analysis

We compared awakening probability across sound intensities for NREM sleep and REM sleep separately (Fig. 1C), between drugs for the three sound intensities and sleep stages separately (Fig. 1d), and as a function of time elapsed after drug injection (Supplementary Fig. 1a).

In combined arousal threshold & optogenetic experiments, awakening probability was computed separately for NREM sleep and REM sleep, across conditions (“S” / “SL” / “L”), and across sound intensities (67/80 dB SPL). Specific conditions with less than 15 trials per condition (e.g. 67 dB SPL in REM sleep) were excluded from analysis.

To assess true synergism between NE activity and sounds above and beyond their expected independent effects, we calculated the expected cumulative effect of the sound alone and the laser alone as following: %S+(100-%S)*(%L)/100 (where %S=awakenings percentage for sound only and %L=awakenings percentage for laser alone, see horizontal lines in Fig. 5a). We then compared it to the observed awakening probability in “SL” condition via 2 way RM ANOVA (between conditions (S/SL) and laser parameters (5Hz/10Hz/burst) in NREM and REM sleep followed by paired t-test corrected with FDR (Fig. 5 and Supplementary Fig. 2)).

#### EEG and EMG analysis

In order to calculate the time-varying amplitude of the EEG in specific frequency bands (0.5-4 Hz for delta, 5-9 Hz for theta) a notch filter was first applied around 50Hz and then the instantaneous amplitude was obtained by taking the absolute value of the Hilbert transform of the band-passed signal. Amplitude signals were then smoothed by a moving average filter (with 0.5 s window) and normalized using z-score relative to the baseline (2 sec before sound). We then averaged the amplitude after the sound (0-6 sec) and compared between trials followed by awakenings and trials followed by maintained sleep. We analyzed the EMG signal by first normalizing using z-score and calculated the increase in the RMS after the sound (0-6 sec) compared to the RMS of the baseline EMG activity (2 sec before sound) in trials followed by awakenings vs. trials followed by maintained sleep. We calculated the FFT of baseline EEG activity for each trial and found decrease in delta power (0.5-4Hz) in trials followed by awakenings compared to trials followed by maintained sleep. These comparisons were done via one-tail paired t-test, for n=9 rats (3 from ChR2-mcherry, 3 from mCherry and 3 from stGtACR2-fRed). We also calculated the FFT of baseline EEG activity for the silencing chronic experiment and found increase in delta power (0.5-4Hz) in trials with laser compared to trials without laser. These comparisons were done via one-tail paired t-test, for n=17 sessions (6 rats injected with stGtACR2-fRed).

#### Sleep spindles

Individual spindles were detected automatically in the EEG as in ^37^. Briefly, we identified events > 2SD over the mean of the 10-16Hz band-pass filtered signal and verified that increased power was specific to this range and not broadband. We then counted the number of spindle peaks that occurred in the 1.5s before sound onset and compared “laser on” trials to “laser off” trials via paired t-test.

#### Locomotor activity analysis

Rat position and movement were analyzed with a custom video software written in Matlab that determines the animal’s center of mass and movement based on subtraction of consecutive images. After initial input from the user (who marks the initial animal location via a dedicated GUI) the software subtracts consecutive frames and, based on the number of pixels that were changed, estimates the degree of movement. Position and movements were compared between saline and yohimbine experiments via paired t-tests (Supplementary Fig. 1d). We then calculated and compared the increase in locomotion time and distance travelled.

#### Pupillometry

To extract pupil area, video images were first cropped around the eye. A mask based on the median values was applied and the best centrally fitted circle was selected using the “regionprops” function in Matlab. For each trial separately, pupil area was normalized by the average baseline pupil area in the 1s preceding trial onset, and percent change dynamics were calculated for the [-5 25]s interval around laser stimulation. Trials were then averaged for each animal and condition separately (∼15 trials per condition) to generate time-courses as seen in Fig. 4b. Statistically significant changes in pupil area in relation to baseline and between parameters conditions (e.g. 70ms vs 90ms pulse duration) were evaluated via Wilcoxon signed-ranked test. The peak (trough) in pupil size was found by searching for the maximal (or minimal) value in the pupil-size vector (excitation or silencing experiments respectively) between 1s before laser offset and 4s after laser offset.

#### Unit identification and spike sorting

Neuronal units were identified using the ‘wave_clus’ software package ^105^ as described previously. For acute activation recordings: extracellular recordings were high-pass filtered above 300 Hz and then a threshold at 5 s.d. above the median noise level was computed (for chronic recordings, a similar threshold was set manually). Then detected events were clustered using superparamagnetic clustering and were categorized as noise or single- or multiunit-clusters. For acute inhibition recordings, units were identified using the KiloSort clustering package ^106^. The Python based clustering package Phy [https://github.com/kwikteam/phy-contrib)] was then used to manually curate Kilosort output in order to select only well isolated single units with clear refractory periods, and to remove artefacts.

#### Analysis of spiking activity

For analysis of spontaneous (Fig. 2d) and sound-evoked (Fig. 2b,e) LC unit activity, data were first divided into vigilances states (active quiet, quiet wake, NREM sleep, NREM-REM sleep transitions, REM sleep) according to EEG, EMG and video (see above). Baseline tonic activity was calculated by averaging the firing rate 2s before sound onset for each unit, and compared between states and within NREM sleep depending on whether the sound led to an awakening or not (Fig. 2b, c). Sound-evoked firing was calculated by averaging the firing rate during the 100ms after sound onset for each unit, and compared between states (Fig. 2g). In addition, the sound-evoked firing rate was divided by baseline tonic activity for each unit separately, and this ratio was compared between states (Fig. 2h). All comparisons were done via a paired t-test.

