## Supplementary for "Locus-coeruleus norepinephrine activity gates sensory-evoked awakenings from sleep"

### Supplementary Figures and Legends

#### Supplementary Figure 1. Effect of NE drugs on sleep architecture and locomotor activity

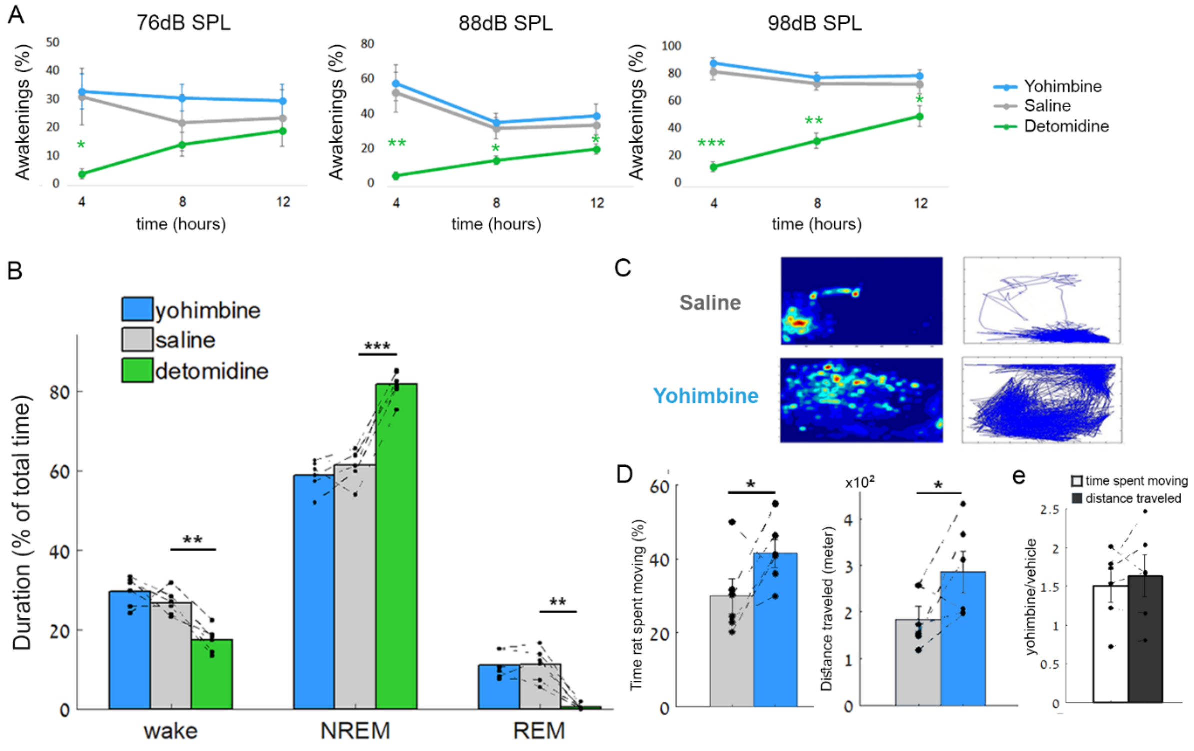


(A) Probability of awakening in response to auditory stimulation as a function of time after drug injection (4/8/12h) for 76 dB SPL (left), 88 dB SPL (middle), and 98 dB SPL (right) sound intensities. Green, detomidine (α2-agonist, 1mg/kg, lower NE); Blue, yohimbine (α2-antagonist, 1mg/kg, higher NE); Gray, saline. Two-way RM ANOVA between drugs and time followed by post-hoc t-test corrected with FDR. Significant decreases in awakening probability are marked by an asterisk (green for detomidine) for specific time intervals (4h/8h/12h).(B) Percent time spent in wakefulness, NREM sleep, and REM sleep during the 12h following drug injection. Colors as above. Two-way RM ANOVA followed by post-hoc t-test corrected with FDR. Note the significant increase in NREM sleep duration for detomidine. (C) Representative heat-map of animal position (left) and tracks showing animal locomotion during the first 3h (right) after injection of saline (top) or yohimbine (bottom). (D) Time spent moving (left), and distance travelled (right) during the first 3h after administration of yohimbine (blue) or saline (gray). (E) Increase in time spent moving (gray, left) and increase in distance travelled (black, right) after yohimbine administration relative to saline reveals that greater locomotor activity goes beyond increase in time spent awake. In all panels data represent mean ± SEM across 6 animals. ***p<0.001, **p<0.01, *p<0.05.

**Supplementary Figure 2. LC opto-activation at multiple stimulation frequencies increases awakening probability**

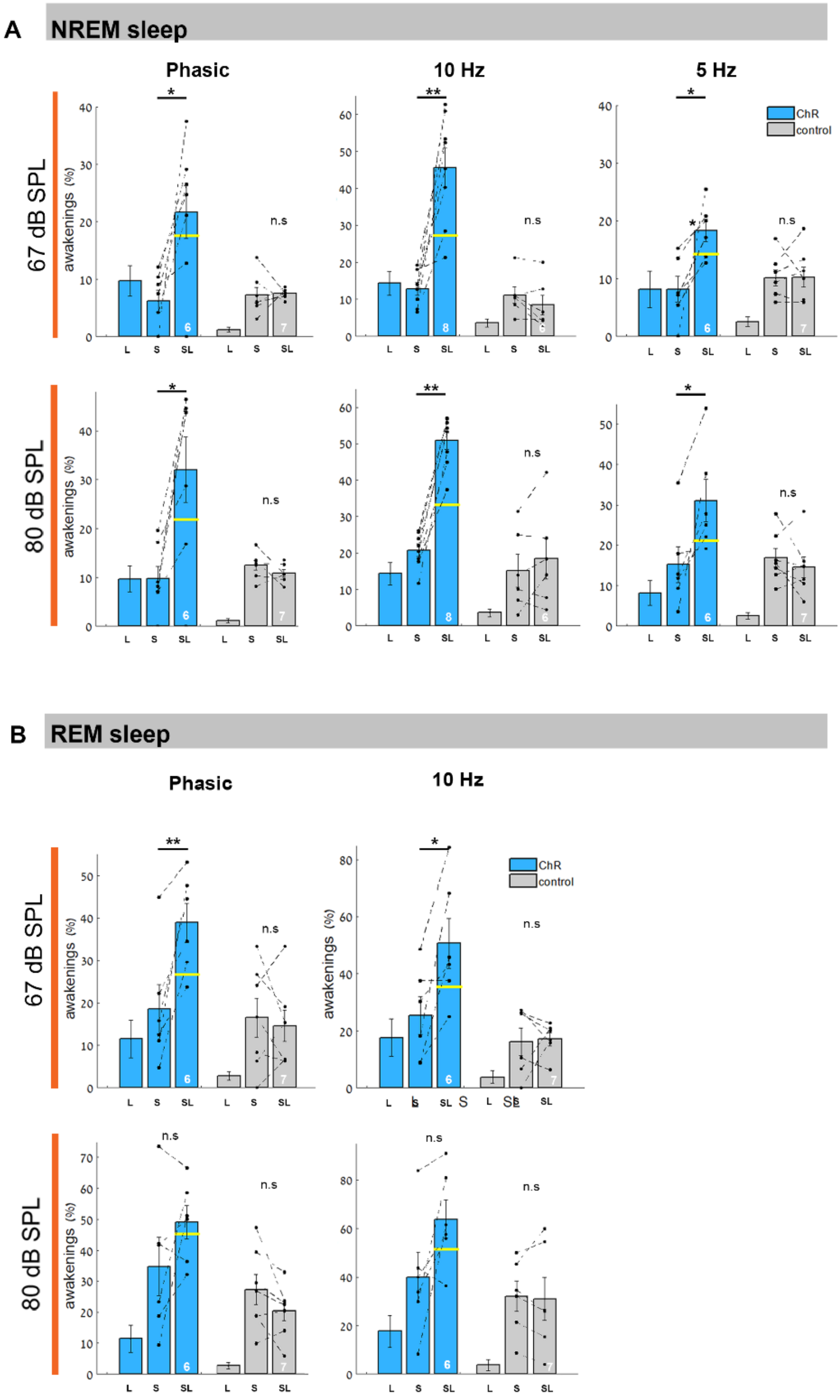

(A) Probability of awakening from NREM sleep following sound presentation (S), sound presented on a background of LC opto-activation (SL), or opto-activation alone (L). Top and bottom rows show data for medium (67 dB SPL) and loud (80 dB SPL) sound intensities, respectively. Blue and gray bars mark data for rats expressing either ChR2-mCherry or mCherry-only controls, respectively. Columns (left to right) show awakening probabilities for these laser stimulation protocols: 'phasic' (burst) stimulation of 3s duration with 10 pulses train at 20Hz repeated at 1Hz (left), 3s duration at 10Hz (middle) or 3s duration at 5Hz (right). Yellow horizontal lines represent the expected independent effect of sound and laser stimulation (Methods). (B) Same as (A) in REM sleep. Two-way RM ANOVA (between conditions and laser parameters) followed by post-hoc t-test corrected with FDR **(***p<0.05, **p<0.01). White numbers on bars mark number of animals contributing to each result.

**Supplementary Fig 3. EEG and EMG associated with awakenings and maintained sleep.**

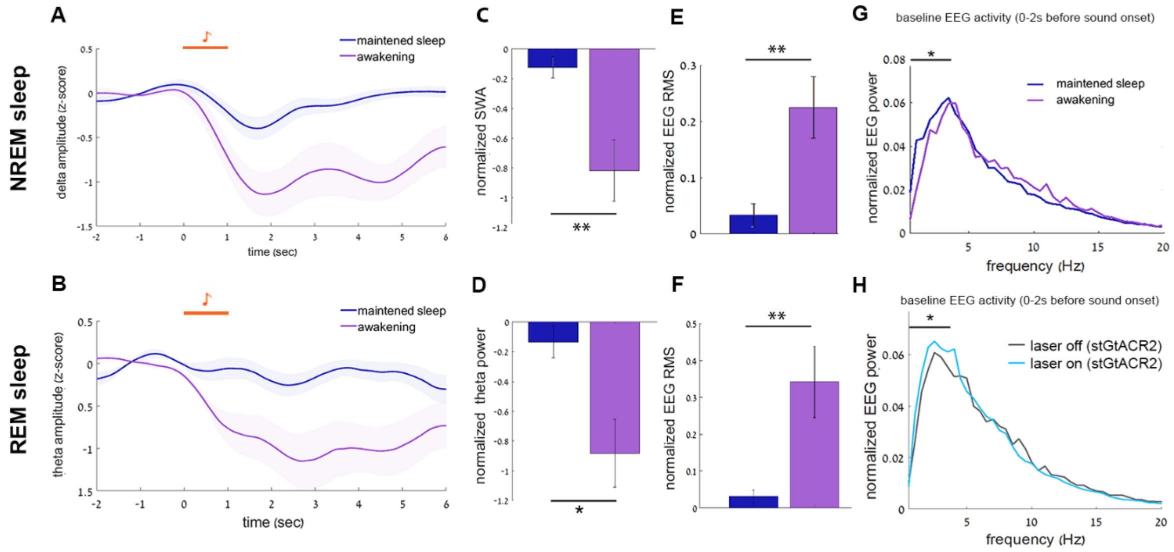


(A,C) Significant decrease in slow wave (0.5–4Hz) amplitude after awakenings from NREM sleep (purple) compared to maintained sleep (blue). (B,D) significant decrease in theta (5-9 Hz) amplitude after awakenings from REM sleep compared to maintained sleep. (E) Significant increase in EMG RMS after awakenings from NREM sleep and (F) from REM sleep, compared to maintained sleep. (G) Normalized EEG power of baseline activity (0-2s before sound onset) when sounds were followed by awakenings (purple) or maintained sleep (blue). Note increase in slow wave (0.5-4 Hz) power for trials followed by maintained sleep (comparisons via one-tail paired t-test, for n=9 (3 rats from ChR2-mcherry, 3 from mCherry and 3 from stGtACR2-fRed). (H) Normalized EEG power of baseline activity (0-2s before sound onset) when sounds were preceded by stGtACR2-fRed laser stimulation (laser on, cyan) or without stimulation (laser off, black). Note increase in slow wave (0.5-4 Hz) power upon LC silencing (n=17 sessions from 6 rats, $p<0.05$ via one-tail paired t-test)

**Supplementary Figure 4. Expression of the control virus (double injection of CAV-PRS.cre together with AAV5-EF1a-DIO-mCherry)**

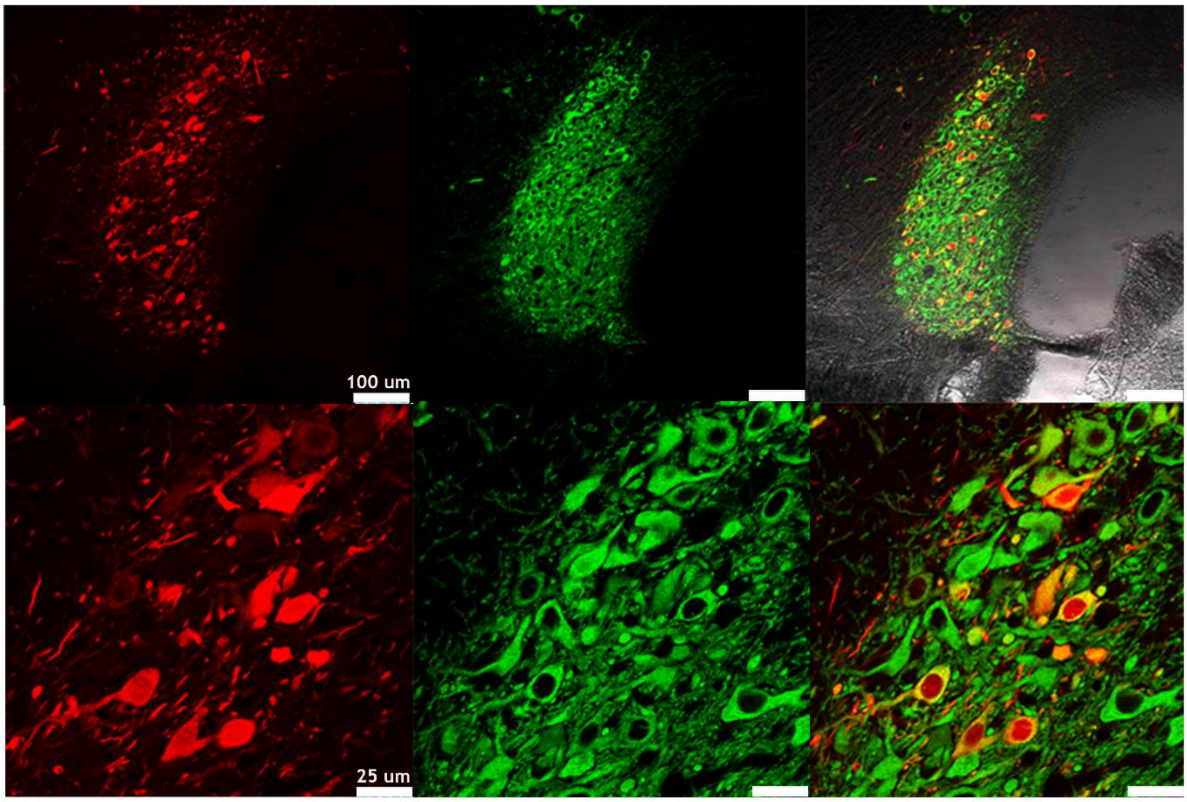


Representative coronal images showing mCherry expression (left column, red), labeling of TH+ neurons (middle column, green), and their overlay (right column, yellow).

**Supplementary Figure 5. Transduction of LC neurons with stGtACR2 introduces a potent opto-activatable, silencing anion conductance.**

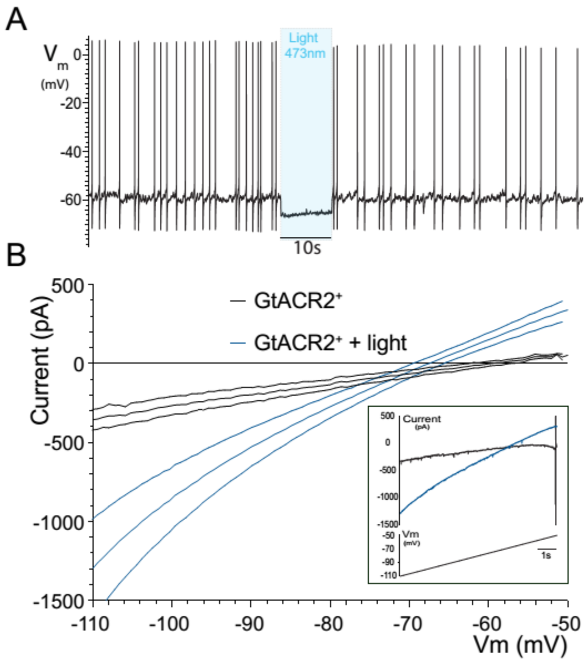


(A) Representative recording from LC neuron showing characteristic spontaneous activity that is silenced by illumination with blue light (473nm, 10s). The resting membrane is rapidly hyperpolarized towards the calculated reversal potential of chloride (-68.7mV). (B) Voltage clamp recording showing the effect of continuous light illumination (473nm x 4mW) on the whole cell current response to a depolarizing ramp (-110 to -50mV over 10 seconds). Inset shows the raw data from the cell in (a) (note action potential discharge close to -50mV prevented by light). Graph shows the V-I relationship LC neurons (dashed line – SEM). Activation of stGtACR2 increased the cell conductance by 336% from 6.3±0.75nS to 21.2±1.46nS (mean ±SEM, n=8). The conductance reversed at -70.9 mV close to the calculated reversal potential for chloride.

**Supplementary Video 1. LC optogenetic excitation leads to pupil dilation**

Optogenetic LC  activation (10Hz, 90ms pulse duration for 3sec, 15mW) under light anesthesia (1.5% isoflurane).
